# Z-chromosome dosage compensation and sex-specific long non-coding RNAs in octopus

**DOI:** 10.1101/2024.12.09.627507

**Authors:** Natali Papanicolaou, Sebastian Wettersten, Alexander Kloosterman, Eduardo Almansa, Eve Seuntjens, Björn Reinius

## Abstract

Sex chromosomes typically present as two copies of a large, gene-rich differentiated chromosome in the homogametic sex (e.g. XX in female mammals, ZZ in male birds) but as a single copy in the heterogametic sex, often paired with a sex-specific, degenerated chromosome (XY in male mammals, ZW in female birds). This creates a significant gene product dosage imbalance of the X and Z chromosomes between the sexes as well as between the single sex chromosome and the autosomes in the heterogametic sex, making its resolution essential. Dosage compensation and its molecular mechanisms have primarily been studied in select organisms, such as primates, rodents, birds, fruit flies, and *Caenorhabditis elegans*, while the extent and mechanisms of dosage compensation remain largely unexplored in the vast majority of animal taxa.

Recently, coleoid cephalopods, including octopuses, squids, and cuttlefish, were reported to possess a ZZ/Z0 sex chromosome system^1^, where males carry two copies of the Z chromosome while females carry only one. However, whether dosage compensation occurs in these species remains unexplored. Here, we show, for the first time, that Z-chromosome dosage compensation is achieved in octopus. Using both original RNA-seq data from *O*.*vulgaris* paralarvae as well as publicly available datasets from *O*.*vulgaris* and *O*.*sinensis*, we report extensive but incomplete dosage compensation of the Z-chromosome in female octopus at the RNA level, similar to Z-chromosome dosage compensation in avian ZW systems. Furthermore, we identify two evolutionarily conserved Z-linked lncRNAs, one featuring strong male-biased expression patterns, that we termed “*Zmast*” and another with strong female-biased expression patterns that we termed “*Zfest*”. Our results provide the first evidence of sex chromosome dosage compensation in octopus, representing one of the most ancient known animal sex-chromosome systems, and raise the possibility that non-coding RNA may play a role in its regulation, akin that observed in younger animal taxa.

## Introduction

Genetic sex determination in animals typically depends on specialized sex chromosomes that differ in copy number between male and female cells. This is exemplified by well-characterized systems such as the XX/XY system in mammals and *Drosophila melanogaster*, where female cells carry two large and gene-rich X chromosomes, while male cells have a single X and a degenerated Y. Conversely, the ZZ/ZW system found in birds, some reptiles, and fish mirrors the XX/XY system, with males being the homogametic sex, carrying two large Z chromosomes, and females carrying a single Z chromosome and a degenerated W^2^. Each of these systems present the hemizygous sex (i.e. XY males in mammals, ZW females in birds) with a considerable gene-dosage imbalance, arising from the presence of only a single copy of the non-degenerated chromosome but two copies of each autosome. The significance of this gene-copy imbalance has driven the evolution of various dosage compensation mechanisms, including X-chromosome upregulation in *Drosophila*^3^, X-chromosome inactivation together with X-upregulation in mammals^4^, and Z-chromosome upregulation in birds^5^, aligning with the theoretical propositions originally put forth by Susumu Ohno^3^. Collectively, these examples underscore the importance of resolving gene dosage imbalances across animal species. X-chromosome dosage compensation in eutherians (e.g. mouse and human) remains the most deeply studied dosage compensation system, with the female-specific long non-coding RNA *Xist* identified as a key regulator, functioning as a silencer of one X-chromosome copy in *cis* in female cells.

Recently, a study on coleoid cephalopod genomes and their evolution, uncovered that various species of coleoid cephalopods including octopuses, squids and cuttlefish possess a ZZ/Z0 sex chromosome system with males carrying two copies of the Z chromosome and females carrying only a single copy, with no evidence for the presence of a degenerated W chromosome^1^, creating a significant dosage imbalance between the single Z in females and the pairs of autosomes. Yet, whether this imbalance is resolved through dosage compensating mechanisms in these animals has not been investigated.

Here, we report, for the first time, extensive but incomplete sex chromosome dosage compensation occurring in coleoid cephalopods, using both publicly available RNA-seq data from adult tissues of two octopus species (*Octopus vulgaris* and *Octopus sinensis*) as well as original high-sensitivity RNA-seq data from *Octopus vulgaris* paralarvae. Intriguingly, we additionally identify two distinct Z-linked lncRNAs, one expressed in a male-specific and another in a female-specific manner, suggesting a putative sex-specific role of these transcripts.

## Results

### Evidence for dosage compensation in *Octopus vulgaris* adult tissues

To investigate sex chromosome dosage compensation in octopus, we first analyzed publicly available RNA-sequencing data from sex-matched tissues (Sub-oesophageal mass - SEB, Supra-oesophageal mass - SEM, optic lobe – OL and Anterior Arm – ARM) of *Octopus vulgaris*^4^ (**Methods**). Given the ZZ/Z0 sex chromosome system, we began by calculating gene-wise Male-to-Female (M:F) expression ratios for each chromosome and tissue to identify potential outlier chromosomes. Indeed, in three out of four tissues examined, *O. vulgaris* chr16 displayed markedly increased M:F expression ratios compared to other chromosomes (chrZ gene-wise gene expression M:F ratios: ARM: 1.15, SEB: 1.05. SEM: 1.04, OL: 0.92; p-value < 0.05, Mann-Whitney-U test for SEB, SEM and ARM, p-value > 0.05, Mann-Whitney-U test, for OL) (**Figure 1A**). Shifted M:F ratios observed across tissues imply a sexually dimorphic dosage specifically of chromosome 16, suggesting that this chromosome constitutes the Z chromosome in *Octopus vulgaris*. Importantly, the observed M:F ratios significantly deviated from the expected value of ~2 that would follow in the absence of any dosage compensation in a ZZ/Z0 system. Deviation from this expected M:F ratio can result from various gene-regulatory modes, including increased transcription from the single female Z chromosome^5–7^, inactivation of one Z in males, reduced transcription from the two male Z chromosomes^8^, reduced^9^ or increased^10^ RNA degradation in females and males respectively, or a combination of these modes, as previously observed in other species.

**Figure 1:**
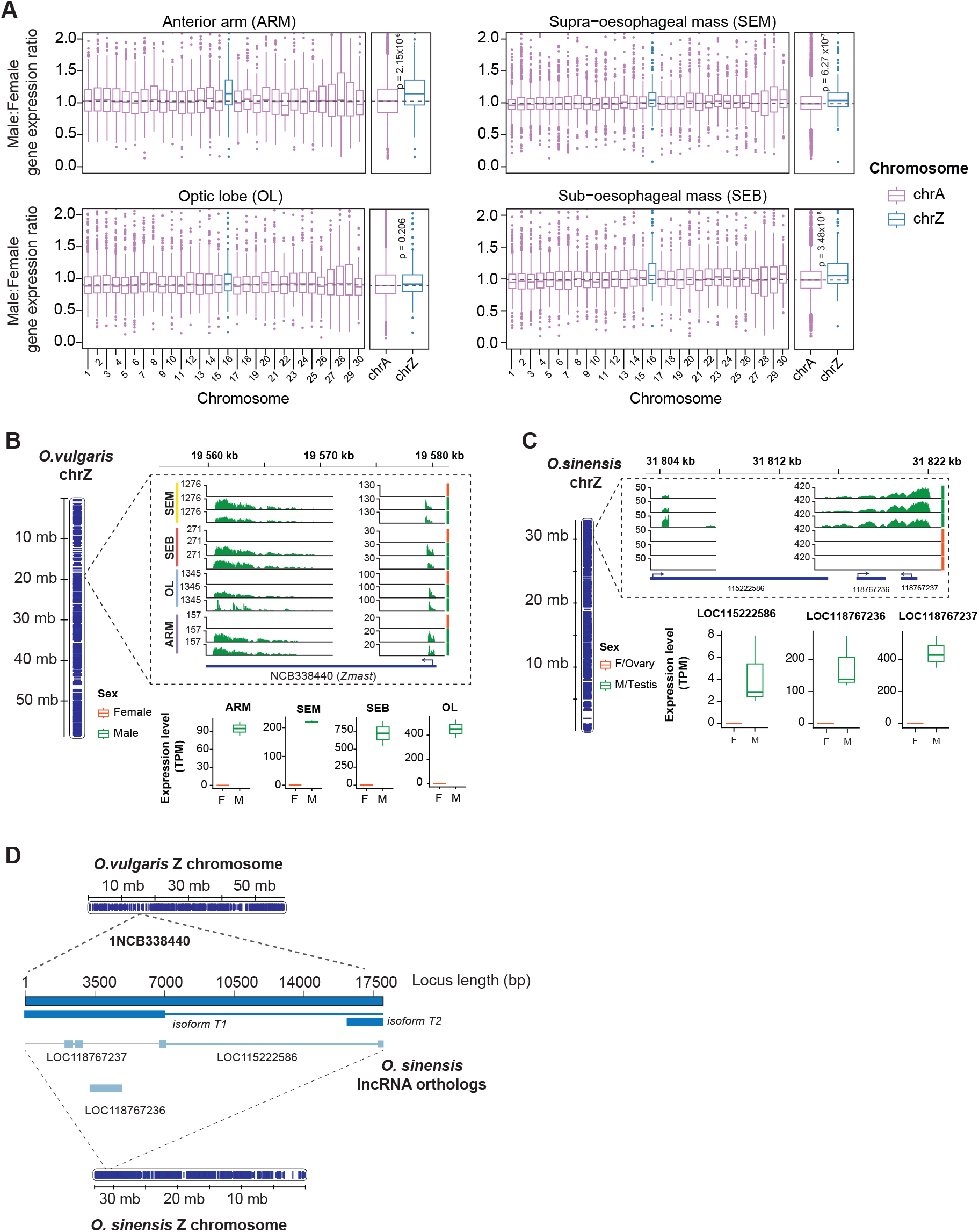
Expression of the *O. vulgaris* Z-chromosome and *Zmast* lncRNA in males and females. **A**. Boxplots of Male:Female gene expression ratios per chromosome and tissue in *Octopus vulgaris*, calculated for expressed genes (TPM > 1) and based on n=2 male and n=1 female samples. Next to each tissue panel, boxplots of Male-to-Female ratios are shown summarised as autosomal (chrA: ARM: n=8376 genes, OL: n=12350 genes, SEM: n=12562 genes, SEB: n=12240 genes) or Z-chromosomes (chrZ: ARM: n=153 genes, OL: n=217 genes, SEM: n=221 genes, SEB: n=212 genes). Median autosomal M:F ratio represented as a dashed line in grey. Autosomal chromosomes shown in purple, and Z-chromosome shown in blue. Data shown as median (center line), first and third quartiles (box limits) and 1.5x interquartile range (whiskers). Mann-Whitney-U test was used for significance testing. **B**. Expression of *Zmast* lncRNA in O. vulgaris tissues. Left: *O. vulgaris* chrZ with length shown in Mb. Right top: Genomic tracks of RNA-seq reads across *Zmast* locus with male samples shown in green and female samples shown in orange, organised by tissue (SEM, SEB, OL, ARM). Right bottom: Expression of *Zmast* lncRNA in TPM per tissue and sex. **C**. Expression of *Zmast* ortholog lncRNAs in *O. sinensis* gonadal tissues. Left: *O. sinensis* chrZ with length shown in Mb. Right top: Genomic tracks of RNA-seq reads across *Zmast* ortholog lncRNAs loci with male samples shown in green and female samples in orange. Right bottom: Expression of *Zmast* ortholog lncRNAs in TPM per tissue. **D**. Sequence alignments between *O. vulgaris Zmast* lncRNA and *O. sinensis Zmast* orthologs.

To determine whether *O. vulgaris* chr16 has sequence similarity to known Z chromosomes, which were previously identified in other octopus species^1^, we ran GENESPACE^11^ using publicly available assemblies of *Octopus bimaculoides*^12^ (GCA_001194135.2) and *Octopus sinensis*^13^ (GCA_006345805.1). We observed a near-perfect conservation of syntenic blocks between *O. vulgaris* chr16 and *O. sinensis* chr LG20 (**Supplementary Figure 1A**), the latter of which was recently identified as a Z chromosome^1^, thereby validating *O. vulgaris* chr16 as the Z chromosome. Interestingly, we observed a similar degree of synteny between *O. vulgaris* chr16 and *O. bimaculoides* chr16, but not chr17 as recently suggested in Coffing et al^1^. We believe that this is a naming discrepancy, possibly due to a large number of unplaced scaffolds in the publicly accessible assembly version of the *O. bimaculoides* genome (ASM119413v2) and not due to disagreement between the syntenic alignments. Based on these findings, we proceeded to compare chromosome expression ratios between the Z chromosome and autosomes in *O. vulgaris*. As female octopuses are expected to be hemizygous for chromosome Z, in absence of dosage compensation, its expression is expected to be considerably lower than those of autosomal pairs of similar gene content. Interestingly, we observed no significant difference between Z-chromosome-expression and autosomal expression in any female tissue examined (median Z-to-autosome: ARM_Female_: 0.83, ARM_Male_: 0.90-0.92, OL_Female_: 0.90, OL_Male_: 0.89-0.93, SEM_Female_: 0.89, SEM_Male_: 0.96-1, SEB_Female_: 1.08, SEB_Male_: 1.27-1.28, p-value > 0.05, Mann-Whitney-U test, **Supplementary Figure 1B**), providing evidence towards increased transcriptional output from the single female Z chromosome to achieve expression levels matching those of autosomes, similar to X-upregulation in mammals^6,7,9,14,15^ and Z-upregulation in birds^5^.

### An evolutionarily conserved Z-linked lncRNA displays male-specific expression in octopus

We proceeded to uncover whether any genes on *O. vulgaris’* putative Z displayed distinct expression signatures between male and female samples. Strikingly, we observed that a lncRNA, *OCTVUL_1NCB338440*, encoded by the Z chromosome, showed high, male-specific, expression in all tissues examined (**Figure 1B**)—thus we called it **Z**-**ma**le-**s**pecific **t**ranscript (*Zmast*). Coding potential calculation^16^ indeed classified this transcript as “*noncoding (weak)*” (alike *Xist*). *Zmast* is encoded in an 18kb-long locus, located between 19.5589–19.5769 Mb, annotated to be transcribed into two isoforms, T1 (18007 nt-long, 19.5590-19.5770, containing one exon (7098 nt-long; chrZ: 19.559– 19.566Mb) and T2 (1969 nt-long; chrZ: 19.575–19.577 Mb), containing two overlapping exons (exon 1: 19.575004–19.576972 Mb; exon 2: 19.576719– 19.576972 Mb) based on current assembly. Using BLAST, we found that a portion of the *Zmast* lncRNA is conserved in three segments, annotated as *LOC115222586, LOC118767237* and *LOC118767236*, in *O. sinensis*, an octopus species that is believed to have diverged from *O. vulgaris* approximately 2.5MYA^17^ (**Figure 2C-D**). To explore the potential male-specific expression of *Zmast* transcripts in this species, we obtained publicly available datasets of RNA-sequencing of gonadal tissue (testis and ovary tissues) from *O. sinensis*^18^. Interestingly, analysis of RNA-sequencing reads and splice junctions suggested that these segments may belong to one and the same transcript (**Supplementary Figure 2A-B**). Indeed, we observed high expression in all three orthologous segments in testis, and the lack thereof in ovaries (**Supplementary Figure 2C**), pointing to an evolutionarily conserved sexually dimorphic expression pattern of this lncRNA. Although the transcriptomic profiles of female and male gonads are expected to be dissimilar and therefore imperfect for calculating M:F ratios, we nonetheless calculated such ratios for the gonads a did observe a tendency for increased M:F ratios for the Z chromosome (LG20) (p=0.0001, Mann-Whitney-U test, **Supplementary Figure 2C**). Using Z-to-autosome (Z:A) ratios calculated within tissues, we observed that in both male and female samples, the transcriptional output of Z was comparable to autosomes (Z:A ratio: Females: 0.64-1.01, Males: 0.93-1.01; p > 0.05, Mann-Whitney-U test), pointing to dosage compensation between the single female Z and the pairs of autosomes (**Supplementary Figure 2D**) similar to our previous observations in *O. vulgaris* tissues.

**Figure 2:**
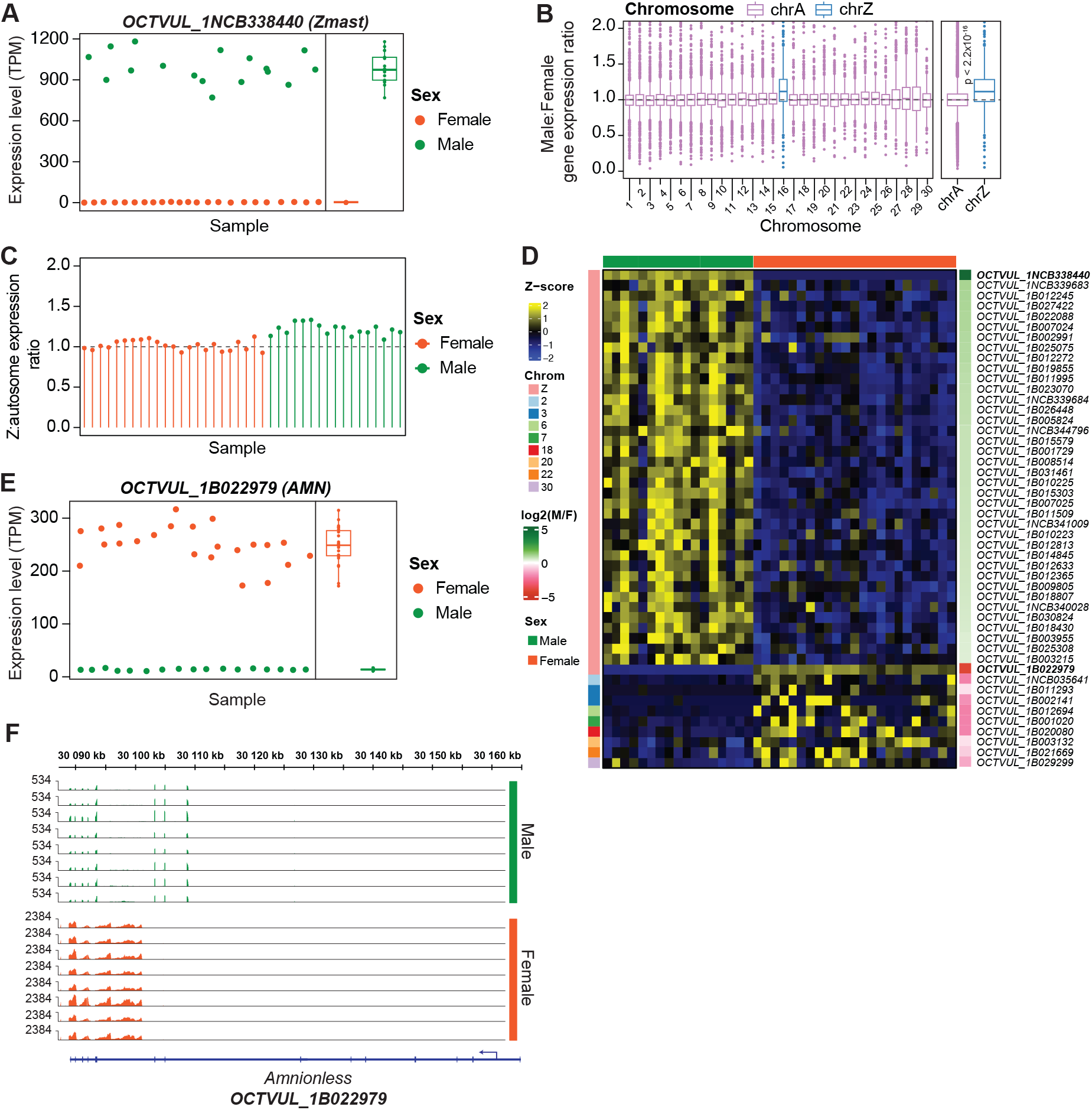
Dosage compensation of the Z-chromosome and *Zfest* expression in *O. vulgaris* paralarvae. **A**. Dot plot and boxplot of Zmast lncRNA expression in *O. vulgaris* paralarvae in TPM based on bulk Smart-seq2 data. Each dot represents an individual sample (total n=40 samples). Male samples (n=17) shown in green and female samples (n=23) shown in orange. **B**. Left: Boxplot of Male:Female gene expression ratios per chromosome in *O. vulgaris* paralarvae, calculated for expressed genes (TPM > 1. Number of genes per chromosome: n = 299 – 1322. Right: Boxplot of M:F gene expression ratio shown summarized for autosomes (n = 19517 total autosomal genes) and the Z chromosome (n = 369). Median autosomal M:F expression ratio shown as a dashed grey line. Autosomal and Z chromosomes shown in purple and blue respectively. Data shown as median (center line), first and third quartiles (box limits) and 1.5x interquartile range (whiskers). Mann-Whitney-U test used for significance testing. **C**. Line and dot plot of median Z-to-autosome expression ratios for each individual sample, calculated for expressed genes (TPM > 1), based on n = 258-273 Z-linked genes and n = 14673-14816 autosomal genes. Genes beyond the 95th percentile of expression per chromosome are excluded. Female samples (n=23) shown in orange and male samples (n=17) shown in green. **D**. Heatmap of z-scores of gene expression for 48 significantly differen-tially expressed genes between male and female *O. vulgaris* paralarvae based on bulk Smart-seq2 RNA-seq data (absolute log2 fold-change > 0.5 and FDR < 0.05), with rows representing genes and columns representing samples. Genes are stratified by chromosome and by male-to-female expression ratios (in log2-scale). Sex represented by top bar, with male samples shown in green and female samples in orange. Left annotation bar shows which chromosome each gene (row) is encoded by. Right annotation bar shows the log2 Male:Female ratio for each gene on the heatmap. Two genes in bold: *OCTVUL_1NCB338440* and *OCTVUL_1B0022979* correspond to lncRNAs *Zmast* and *Zfest*, respectively. **E**. Dot plot and boxplot of *OCTVUL_1B022979* (*Amnionless*) expression in *O. vulgaris* paralarvae in TPM based on bulk Smart-seq2 data. Each dot represents an individual sample. Male samples shown in green and female samples shown in orange. **F**. Genomic coverage tracks of RNA-seq reads based on *O. vulgaris* paralarvae bulk Smart-seq2 RNA-seq data over the *Amnionless* locus (*OCTVUL_1B022979*) shown for 8 representative male samples (denoted by green bar on the right side of the panel) and 8 representative female samples (denoted by the orange bar on the right side of the panel).

### Z-chromosome upregulation and a female-specific lncRNA in paralarvae

To independently validate and more extensively characterize Z-chromosome dosage compensation, we bred and collected 40 three-day old *Octopus vulgaris* paralarvae and prepared mini-bulk Smart-seq2 libraries from 3 day old *Octopus vulgaris* paralarvae, obtaining full-transcript length and high sensitivity^19^ **(Methods)**. We detected an average 20724 genes across the 40 samples (TPM > 1), 398 of which were Z-linked (**Supplementary Figure 2A-B**). Confirming our previous observations in *O. vulgaris* adult tissues, *Zmast* displayed a binary pattern of expression with extremely high expression in 17 samples, which we identified as male (mean TPM = 990 ± 112.4, n = 17; range: 771.0– 1182.0) and low expression in 23 samples, which we identified as female (mean TPM = 3.8 ± 1.42, n=23; range: 1.18-6.6, **Figure 2A**), in line with the expected rate of male and female samples from random embryo sampling. Having identified the sex of the paralarvae samples, we calculated M:F Z-gene expression ratios and confirmed a significant, incomplete degree of dosage rebalancing of the *O*.*vulgaris* Z-chromosome (gene-wise M:F expression ratio: 1.12, p-value < 2.2×10^−16^, Mann-Whitney-U test, **Figure 2B**). Ratios of Z-chromosome-to-autosome expression levels showed that the female Z-chromosome is transcriptionally upregulated compared to autosomal chromosomes (**Figure 2C** and **Supplementary Figure 3C**). Differential expression analysis between the sexes resulted in the identification of 48 differentially expressed genes (0.3% of expressed genes), 39 of which were encoded by the Z-chromosome (**Figure 2D**). These results indicate that overall gene expression patterns are broadly similar between male and female *Octopus vulgaris* paralarvae, and at the same time that sex-biased genes are disproportionately concentrated on the Z chromosome. Notably, 38 out of these 39 Z-genes showed a male-biased expression level. The magnitude of the male-bias in 37 out of the 38 genes were in the range of 1.41 – 2-fold, and therefore in the expected range resulting from incomplete dosage compensation of the Z chromosome, while *Zmast* was identified as truly male-specific (256-fold increase compared to females). The single remaining Z-linked gene, *OCTVUL_1B022979*, representing the gene *Amnionless* (*AMN*), presented a strongly female-biased expression in all samples examined (18-fold increase compared to males), mirroring the pattern of *Zmast*. Investigating the sex-biased expression pattern of *AMN* more closely, we found that, while male samples showed the expected exon-level expression, consistent with annotated exons and splice junctions, female samples did not, displaying instead a distinct pattern of expression, spanning both annotated intronic and exonic regions of the *AMN* gene **(Figure 2E)**. This pattern suggests a potentially anti-sense transcript produced from the *AMN* locus and only expressed in females (**Figure 2F**). Using RNA-seq data from *O. vulgaris* adult tissues^4^, we confirmed the expression of the distinct 13.8 kb female-specific transcript (**Supplementary Figure 4A**), demonstrating that its expression is not specific to the to the paralarval stage. This suggests it may serve a more fundamental role across multiple tissues and developmental stages. We next sought to examine whether this transcript shows evolutionary conservation and similar expression patterns in *O. sinensis* (East Asian octopus). Strikingly, we found that the *AMN* locus in *O. sinensis* not only showed a similar pattern of expression as in *O. vulgaris*, but that the female-specific expression pattern indeed corresponds to an annotated antisense lncRNA, *LOC118767298* (**Supplementary Figure 4B**). Given its heavily female-biased expression pattern, we termed this lncRNA, *Zfest*, for **Z**-**fe**male-**s**pecific **t**ranscript.

## Discussion

The recent revelation of a ZZ/Z0 sex chromosome system in coleoid cephalopods including octopus, cuttlefish, squid, and the chambered nautilus by Coffing and colleagues marks a landmark discovery that pushed the evolutionary age of animal sex chromosome systems to ~480MYA^1^. The revealed hemizygosity begged the question of whether the Z-dosage discrepancy in female cephalopod cells is resolved at the transcriptional level similarly to other sex chromosome systems investigated across the animal kingdom. Using RNA-sequencing datasets from matched *O. vulgaris* somatic tissues^4^ and *O. sine is*^18^ adult gonadal tissues, we found that Male-to-Female ratios of Z-chromosome gene expression were significantly lower than 2, which would be the expected ratio in the absence of any dosage-compensating mechanism, while being significantly above 1.0, thereby indicating partial dosage compensation. By generating original bulk Smart-seq2 data from *O. vulgaris* paralarvae, we confirmed our findings of extensive, yet incomplete, transcriptional-level Z-chromosome dosage compensation, an observation which bears striking similarity to the partial Z-chromosome dosage compensation found in the Z/W-system of birds^5^. Furthermore, we observed near-balanced Z-to-autosome ratios within samples, indicating that the transcriptional output of the single female Z chromosome is upregulated to match the expression of the two autosomes, something that was previously observed in mammals^6,7,15^, birds^5^, *Drosophila melanogaster*^20,21^ and *Caenorhabditis elegans*^14^, extending this apparently fundamental mode of dosage compensation to coleoid cephalopods.. Intriguingly, we identified a lncRNA, “*Zmast”* (*OCTVUL_1NCB338440*), located on *O. vulgaris* chromosome Z (annotated as chr16 in GCA_951406725.2 genome assembly), displaying high male-specific expression, which is conserved in *O. sinensis* in three shorter annotated and neighboring non-coding sequences (*LOC115222586, LOC118767237* and *LOC118767236*) and exclusively expressed in males also in this species. Our current data suggest that these annotated segments represent the same transcribed unit. We also identified a second Z-linked lncRNA, 13.8 kb in length, that is strongly female-biased in its expression, which we termed *“Zfest”*, and which is transcribed from the *Amnionless* locus in *O. vulgaris*, conserved in *O. sinensis*, and annotated as a lncRNA transcribed in the antisense direction relative to the *Amnionless* host gene. LncRNAs have long been implicated as master regulators of sex-chromosome dosage compensation in various species and sex-chromosome systems, such as *roX1* and *roX2* responsible for X-chromosome upregulation in Drosophila^21^, and *Xist*^22,23^ and *Rsx*^24^, responsible for X-chromosome inactivation in eutherians and marsupials respectively. More recently, two lncRNAs *MAYEX and FEREX*, were shown to display male-specific and female-specific expression patterns respectively in the green anole lizard, with *MAYEX* being responsive for X-chromosome upregulation of the single male X chromosome in males^25^. Given the well-established role of lncRNAs in sex chromosome dosage compensation across diverse species and the strikingly mirrored male/female expression of *Zmast* and *Zfest*, the possibility that these lncRNAs operate in Z-chromosome dosage compensation in cephalopods is particularly intriguing. If one, or both, of these lncRNAs are involved in dosage compensation, they would likely act through distinct mechanisms. *Zmast* could act by either downregulating the transcriptional output of both Z chromosome similar to X-chromosome dampening in human preimplantation development^26^, or, by inactivating one of the two Z chromosomes in the male, alike the mechanism of X-inactivation in mammals^22^. *Zfest* on the other hand could act by orchestrating the transcriptional upregulation of the single female Z chromosome as observed in the green anole lizard^25^ and Drosophila^27^. While the *Zmast* and *Zfest* exhibit sex-specific smoking-gun expression patterns suggestive of lncRNAs involved in dosage compensation, further mechanistic studies are needed to validate their functional roles.

As the most recent estimates suggest that the cephalopod Z chromosome may be the oldest extant sex chromosome, the presence of dosage compensating mechanisms in this system highlights the fundamental need for the evolution of sex chromosome dosage compensating mechanisms across different the animal kingdom.

## Methods

### Ethical considerations

Adult male and female wild caught O. vulgaris were maintained and mated in the Oceanographic Center of the Canary Islands Instituto Espanol de Oceanografia (IEO-CSIC, Tenerife, Spain) according to Almansa et al. Upon spawning and hatching, paralarvae were collected in 200 ul of cold TRI reagent. All procedures were approved by the ethical board and competent authority on animal experimentation from CSIC (permit CEIBA 1377-2013) in compliance with Directive 2010/63/EU.

### Re-analysis of publicly available RNA-seq datasets of *Octopus vulgaris* and *Octopus sinensis* samples

#### FASTQ file pre-processing and STAR mapping

*Octopus vulgaris* and *Octopus sinensis* bulk tissue RNA-seq fastq files^4,18^ were obtained from ENA under accession numbers: E-MTAB-3957 and SRP499429. The latest available assemblies of the *O. vulgaris* and *O. sinensis*^13^ genome was obtained from NCBI (assemblies: GCA_951406725.2, GCA_006345805.1). To improve mapping accuracy, STAR^28^ (v.2.7.2a) indexes for the respective genomes were created prior to mapping using both genome fasta and GTF files (STAR –runMode genomeGenerate, --sjdbGTFfile *.gtf). Raw fastq files were aligned to the respective genomes and transcriptomes and reads were quantified using STAR (v.2.7.2a) with the following options: --runMode alignReads, --clip3pAdapterSeq AGATCGGAAGAGCACACGTCTGAACTCCAGTC A AGATCGGAAGAGCGTCGTGTAGGGAAAGAGTGT --quantMode TranscriptomeSAM GeneCounts, --outSAMtype BAM Unsorted SortedByCoordinate Gene expression read counts were TPM normalized by dividing gene counts by the effective gene length in kb (RPK), dividing the total gene-length-normalized counts by 10^6^ to calculate the scaling factor and dividing the RPK by the scaling factor.

#### Expression ratio calculations

Gene-wise Male:Female expression ratios were calculated for expressed genes (TPM >1) per tissue for *O. vulgaris* and *O. sinensis* data. Chromosome:Autosome expression ratios were calculated per sample, for expressed genes (TPM >1) and after exclusion of the top 5% expressed genes per chromosome and tissue to avoid bias from highly expressed tissue-specific genes.

#### Differential gene expression analysis

Differential gene expression analysis was performed using the DESeq2 software^29^ package (v1.38.3) in R (v4.2.2). Briefly, quantified gene read count tables for *O. vulgaris* samples were obtained from STAR (see section “***Processing of publicly available RNA-seq datasets***.”) and were used as input for DESeq2. To obtain DEGs between the two sexes, DESeq was run for each individual tissue type (ARM, OL, SEM, SEB) separately. Only DEGs with log2FC > 0.5 | < −0.5 were kept.

### Nucleotide BLAST (BLASTN)

Discontiguous megablast was used to compare *O. vulgaris Zmast* lncRNA exonic and intronic sequences to genomic sequences of *O. sinensis* with default settings. Sequences producing significant alignments with percentage identity more than 90% and query cover more than 80% were considered for further analysis.

### Synteny calculation

The latest available assemblies of the *Octopus bimaculoides, Octopus sinensis* and *Octopus vulgaris* genome were obtained from NCBI (assemblies: GCF_001194135.2, GCF_006345805.1 and GCA_951406725.2 respectively). From these assemblies the genomic GFF files and the translated CDS were used for analysis. Analysis was performed using R Statistical Software (v4.1.2) and the R package GENESPACE (v1.3.1)^11^. Default GENESPACE workflow and output was used to plot the syntenic block analysis for all chromosomes. Region of interest syntenic block plot was made using *O. vulgaris* as reference with prior GENESPACE output, without rendering background and using chromosome 16 as the region of interest.

### RNA extraction from *O*.*vulgaris* paralarvae

Octopus vulgaris paralarvae were collected 2 days post-hatching in 200 ul of TRI reagent and placed in −80°C. To improve sample lysis, 200 ul of ice-cold TRI reagent was added to each sample and each sample was well homogenized using a Fisherbrand 150 handheld probe homogenizer with the probe thoroughly washed between samples by activating the homogenizer during submersion in water, followed by 70% ethanol and TRI reagent to ensure effective cleaning and minimize the possibility of cross-contamination between samples. The samples were immediately returned to −80°C. Immediately before extraction, the samples were placed on ice to partially thaw and 600 ul of cold TRI reagent was added to each sample to achieve a final TRI reagent volume of 1ml. RNA extraction was performed according to the manufacturer’s protocol. Briefly, the samples were thawed on ice and placed at room temperature for 5 minutes to allow for dissociation of nucleoprotein complexes, after which 200 ul of chloroform was added to each sample. The samples were then shaken vigorously for 30 seconds and allowed to stand at room temperature for 5 minutes. The resulting mixture was centrifuged at 12000xg for 15 minutes at 4°C. To extract and precipitate the RNA, the aqueous phase was removed and transferred to a fresh Eppendorf tube and topped with 500 ul of isopropanol. The samples were mixed by inversion and incubated at room temperature for 10 minutes to facilitate RNA precipitation. Next, the samples were centrifuged at 12000xg for 10 minutes at 4°C, the supernatant was carefully removed and discarded and the RNA pellet washed by adding 1 ml of freshly-prepared 75% ethanol, brief vortexing and centrifuging for 5 minutes at 7500xg at 4°C. The wash was repeated one more time and the resulting RNA pellet was air-dried for 10 minutes until most of remaining ethanol was evaporated. Finally, the RNA pellet was resuspended in 30 ul of nuclease-free water, incubated at 60°C for 15 minutes and the concentration was measured on a Nanodrop 2000 instrument. The resulting samples were stored in −80°C.

### Mini-bulk Smartseq2 library preparation from

#### *O*.*vulgaris* paralarvae

Mini-bulk Smart-seq2 libraries were prepared as previously described^5,19^. First, total RNA was diluted to 1ng/μL, of which 2 ng (2 μL) was added to 2.5 μL of mini-bulk SS2 lysis buffer containing 1 μL of 10 mM dNTP mix [Roche], 0.11 μL of 50 massunits/μL SEQURNA thermostable RNase inhibitor^30^ [SEQURNA], 1 μL of 10 μM oligodT [IDT; 5’– AAGCAGTGGTATCAACGCAGAGTACT30VN-3’] and 0.39 μL of nuclease-free water [Ambion]. *<u>Reverse transcription and PCR preamplification</u>*. The samples were then incubated at 72°C for 3 minutes and spun down before adding 5.5 μL of reverse transcription mastermix to each sample (2 µL of 5x SuperScript II buffer [Thermo Fisher Scientific], 0.5 µL of 0.1 M DTT [Thermo Fisher Scientific], 2 µL of 5 M betaine solution [Sigma-Aldrich], 0.1 µL of 1 M MgCl_2_ [Ambion], 0.1 µL of 100 µM TSO [IDT; 5’-AAGCAGTGGTATCAACGCAGAGTACATrGrG+G-3’], 0.5 µL of SuperScript II reverse transcriptase [Thermo Fisher Scientific], and 0.3 µL of nuclease-free water [Ambion]). The samples were then briefly centrifuged at 500xg for 10 seconds at room temperature, before starting the reaction in a thermocycler with a heated lid using the following program: 1 cycle of 42°C for 90 minutes, 10 cycles of [50°C for 2 minutes, 42°C for 2 minutes], 1 cycle of 72°C for 15 minutes followed by 4°C on hold. The resulting cDNA was spun down and PCR preamplification was performed by adding 15 μL of PCR mastermix to each sample (PCR mastermix composition: 12.5 μL of 2x KAPA HiFi HotStart readymix [Roche], 0.2 μL of 10 µM ISPCR primers [IDT; 5’-AAGCAGTGGTATCAACGCAGAGT-3’], 2.3 μL of nuclease-free water [Ambion]), spinning down at 500 x g for 10 seconds at room temperature and performing the PCR in a thermocycler with a heated lid using the following program: 1 cycle of 98°C for 3 minutes, 14 cycles of [98°C for 20 seconds, 67°C for 15 seconds, 72°C for 6 minutes], 1 cycle of 72°C for 5 minutes and 4°C on hold. *<u>Bead purification of cDNA libraries</u>*. The resulting cDNA libraries were purified by adding 20 μL of AMPure XP beads [Beckman-Coulter] to each sample, thoroughly mixing by pipetting up and down 10 times and incubating at room temperature for 8 minutes, before placing on a magnetic rack for 5 minutes or until the samples appear clear. The supernatant was carefully removed and discarded and the bead pellets were washed by adding 200 μL of freshly-prepared 80% ethanol, incubating for 30 seconds (while on magnetic rack), removing and discarding the liquid without disturbing the pellet. The ethanol wash was performed once more before letting the beads air-dry for 5 minutes while on magnetic rack. Finally, the beads were resuspended in 18 μL of nuclease-free water (off the magnetic rack), incubated at room temperature for 5 minutes and placed back on the magnetic rack for 2 minutes before collecting 16 uµ of each sample (to avoid bead contamination) and transferred to a new tube. *<u>Quality assessment and library quantification</u>*. Library quality was assessed on an Agilent Tapestation 4200 instrument using a D5000 DNA ScreenTape kit following the manufacturer’s instructions. Library concentration was measured using the QuantiFluor dsDNA kit [Promega] according to the manufacturer’s protocol on a Thermofisher Scientific Varioskan Lux instrument. *<u>Library tagmentation</u>*. To prepare for the tagmentation reaction, 3 ul of each cDNA library was normalized to 500 pg/μL using nuclease-free water [Ambion]. 1 ng of cDNA (2 μL) was then combined with 18 μL of tagmentation mastermix containing 2 μL of 10x TAPS-MgCl_2_ buffer (100 mM TAPS [Sigma-Aldrich], 50 mM MgCl_2_ [Sigma-Aldrich], 5 µl of 40% PEG-8000 solution [Sigma-Aldrich], 10.18 μL nuclease-free water and 0.82 μL 27 µM Tn5 [in-house produced]. The tagmentation reaction was performed in a thermocycler with a heated lid at 55°C for 8 minutes. To strip the Tn5 from the tagmented cDNA, 3.5 μL of 0.2% SDS solution was added to each sample, followed by thorough mixing, centrifugation at 500 x g for 10 seconds and incubation at room temperature for 5 minutes. *<u>PCR amplification of tagmented libraries</u>*. 1.5 μL of combined Nextera i7 and i5 indexes (1 µM each) were added to each sample as well as 16.5 μL of PCR amplification mastermix (10 μL of 5x KAPA HiFi PCR buffer [Roche], 1.5 μL of 10 mM dNTP mix [Roche], 1 μL KAPA HiFi polymerase enzyme [Roche] and 4 μL of nuclease-free water [Ambion]). PCR amplification of the tagmented libraries was performed in a thermocycler with a heated lid using the following program: 1 cycle of 72°C for 3 minutes, 1 cycle of 95°C for 30 seconds, 12 cycles of [95°C for 10 seconds, 55°C for 30 seconds, 72°C for 30 seconds], 1 cycle of 72°C for 5 minutes and 4°C on hold. The libraries were then pooled and bead purification took place as described above. The final pool concentration was measured on a Qubit 4.0 instrument using a Qubit dsDNA High Sensitivity Assay Kit [Molecular Probes] and the library fragment distribution was inspected on an Agilent Tapestation 4200 instrument using a D5000 DNA ScreenTape kit following the manufacturer’s instructions. The library pool was converted using a MGIEasy Universal Library Conversion Kit (App-A) according to the manufacturer’s instructions and sequenced on a MGI DNBSEQ G400-RS instrument using a G400 App-D FCL PE100 kit (Read 1 = 100 bp, Read 2 = 100 bp, Index 1 = 10 bp, Index 2 = 10 bp).

#### Sequencing data pre-processing

Raw MGI fastq files were demultiplexed using mgikit (v1.0.0)^31^, which required the creation of a sample sheet containing sample names and their respective i7 and i5 barcodes. The demultiplexing was done for each lane using mgikit demultiplex (−m 2 --template i710:i510). The demultiplexed samples were then merged across the lanes to create demultiplexed fastq files containing all reads across the four lanes. The mapping and analysis of the data was performed as described above (see section “**Re-analysis of publicly available RNA-seq datasets of *Octopus vulgaris* and *Octopus sinensis* samples**”).

---

**Supplementary Figure 1.**
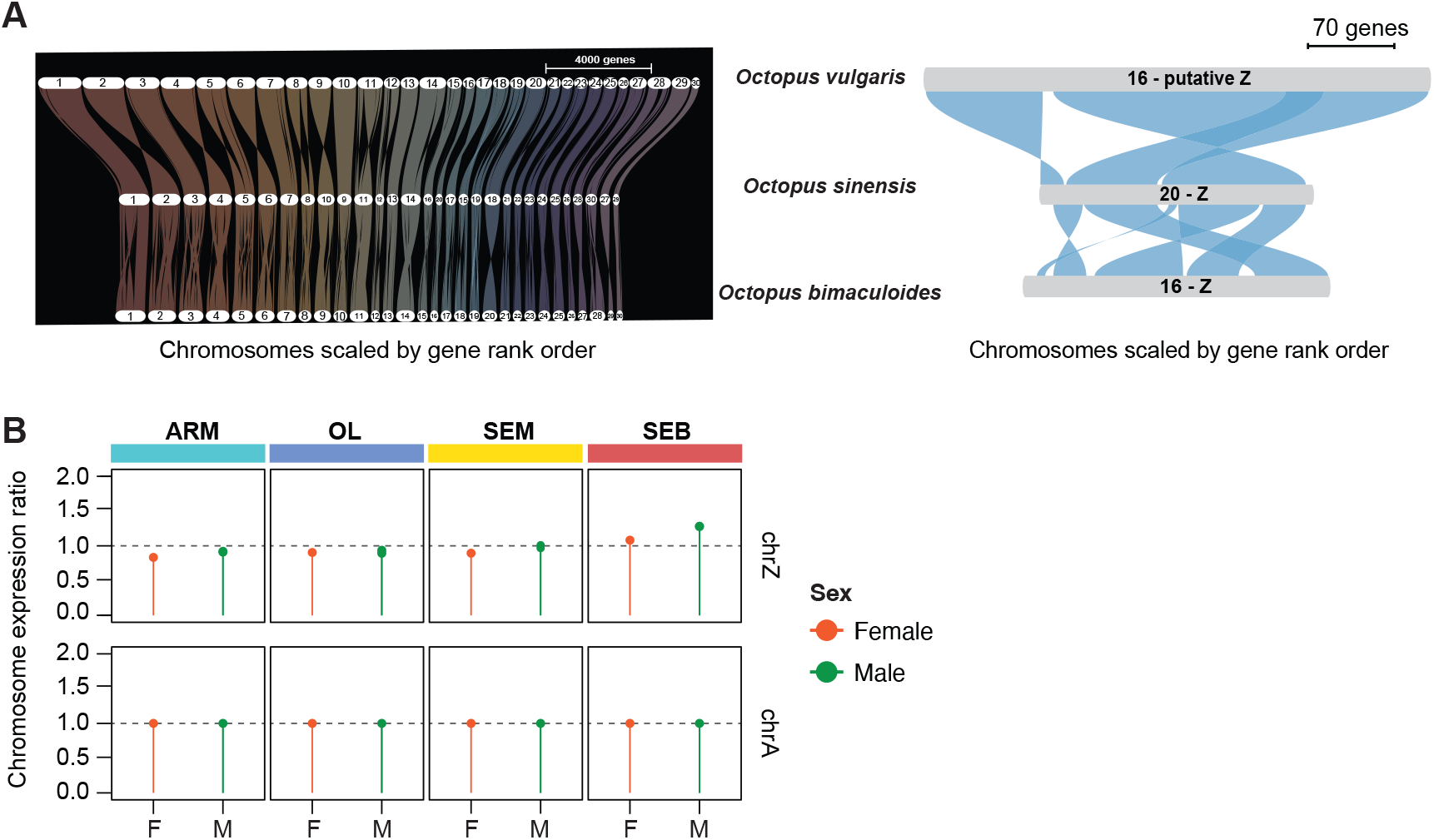
Syntenic alignment of the *O. vulgaris* Z chromosome. **A**. GENESPACE output of chromosomal syntenic alignments between *O. vulgaris, O. sinensis* and *O. bimaculoides* for all 30 chromosomes (left) and for the Z chromosome (right). **B**. Line and dot plot of median chromo-some-to-autosome ratios in *O. vulgaris* tissues for male and female samples with Z-to-autosome ratios shown at the top and autosomal ratios shown at the bottom. Tissue type is denoted with colored bars over individual panels. Only expressed genes (TPM > 1) and genes within the 95th percentile per chromosome and tissue were included (ARM: chrA n = 7539-7616 genes, chrZ: n = 137-146 genes, OL: chrA n = 11396-11497 genes, chrZ n = 201-205 genes, SEM: chrA n = 11598-11633 genes, chrZ n=203-206 genes, SEB: chrA n = 11345-11355 genes, chrZ n = 197-198 genes). Female samples shown in orange and male samples shown in green.

**Supplementary Figure 2.**
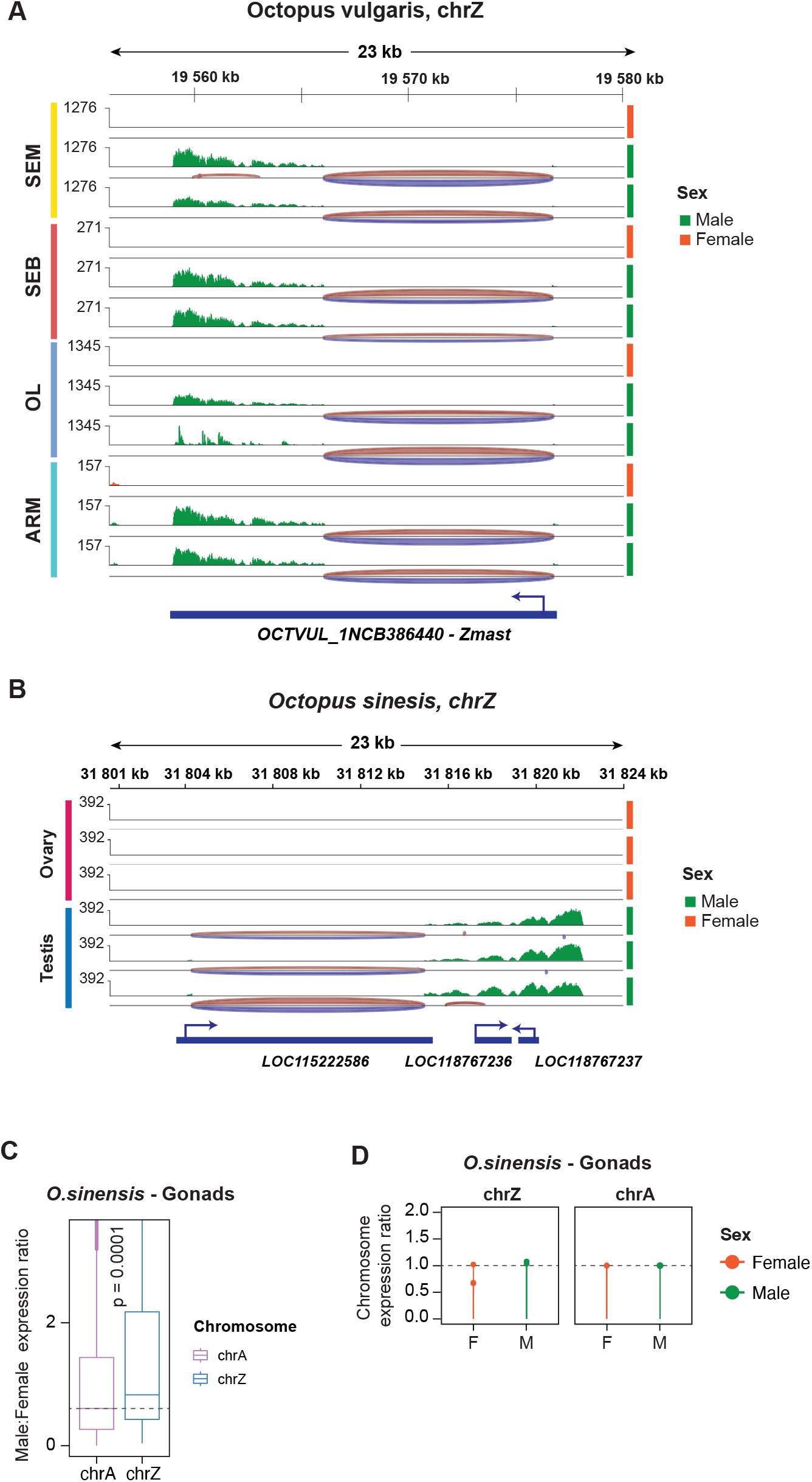
*Zmast* shows similar splicing patterns in *O. vulgaris* and *O. sinensis*. **A**. Genomic coverage tracks of *Zmast* lncRNA expression and splice junctions in *O. vulgaris* tissues for male and female samples based on bulk RNA-seq. Female and male samples are shown in orange and green respectively. Samples were group-autoscaled based on tissue type (SEM, SEB, OL, ARM). **B**. Genomic coverage tracks of *Zmast* ortholog lncRNAs expression and splice junctions in *O. sinensis* gonadal tissues (ovary or testis) based on bulk RNA-seq data. Female and male samples shown in orange and green respectively. **C**. Boxplot of male-to-female gene expression ratio for autosomes (chrA, n = 9013 genes) and the Z chromosome (chrZ, n = 163 genes) based on bulk RNA-seq data from *O. sinensis* gonadal tissues (male testis, female ovary). Autosomes denoted in purple, Z chromosome denoted in blue. Mann-Whitney-U test used for significance testing. Data shown as median (center line), first and third quartiles (box limits) and 1.5x interquartile range (whiskers). **D**. Line and dot plot of median chromosome-to-autosome ratios in *O. sinensis* tissues for male and female samples with Z-to-autosome ratios shown on the left and autosomal ratios shown on the right, based on n= 8424 – 9795 autosomal genes and n=154-168 Z-linked genes. Tissue type is denoted with colored bars over individual panels. Only expressed genes (TPM > 1) and genes within the 95th percentile per chromosome and tissue were included. Female samples shown in orange and male samples shown in green.

**Supplementary Figure 3.**
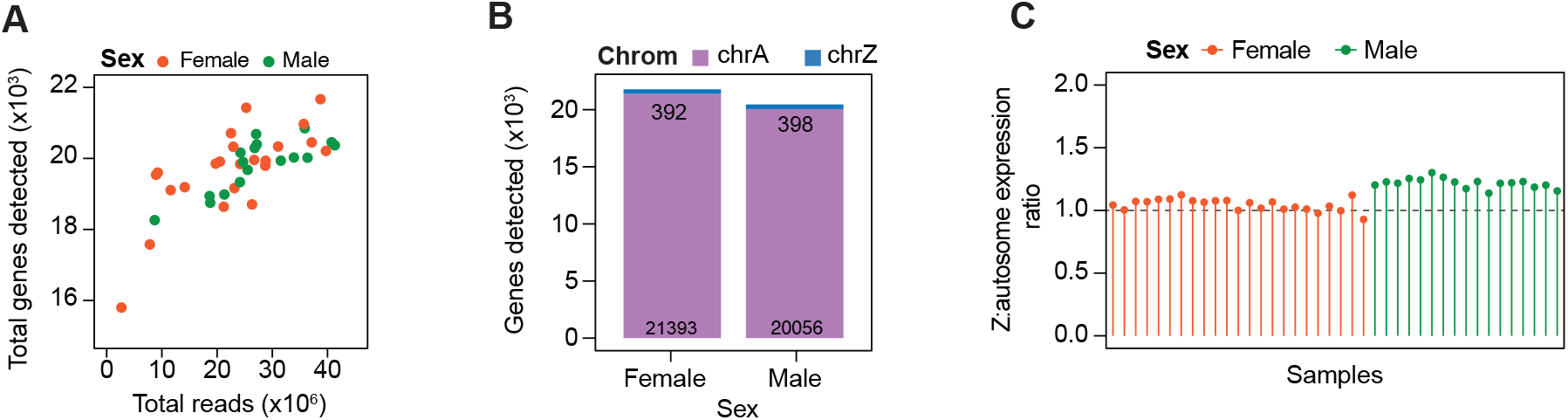
Quality control of bulk Smart-seq2 in *O. vulgaris* paralarvae. **A**. Scatterplot of total number of reads (in millions) on the x-axis versus total number of genes detected (in thousands, TPM > 1) on the y-axis per sample (n = 40) in bulk Smart-seq2 of *O. vulgaris* paralarvae. Female samples shown in orange and male samples shown in green. **B**. Barplot of total number of genes detected (TPM > 1), divided into autosomal genes (in purple) and Z-chromosome genes (in blue) across male and female *O. vulgaris* paralarvae using bulk Smart-seq2 RNA-seq. **C**. Line and dot plot of median Z-to-autosome expression ratio shown for each female (n = 23) and male (n = 17) sample of O. vulgaris paralarvae based on bulk Smart-seq2 RNA-seq data. Similar to Figure 2C, only expressed genes (TPM > 1, n= 290 chrZ genes) were included but without upper gene expression cutoff.

**Supplementary Figure 4.**
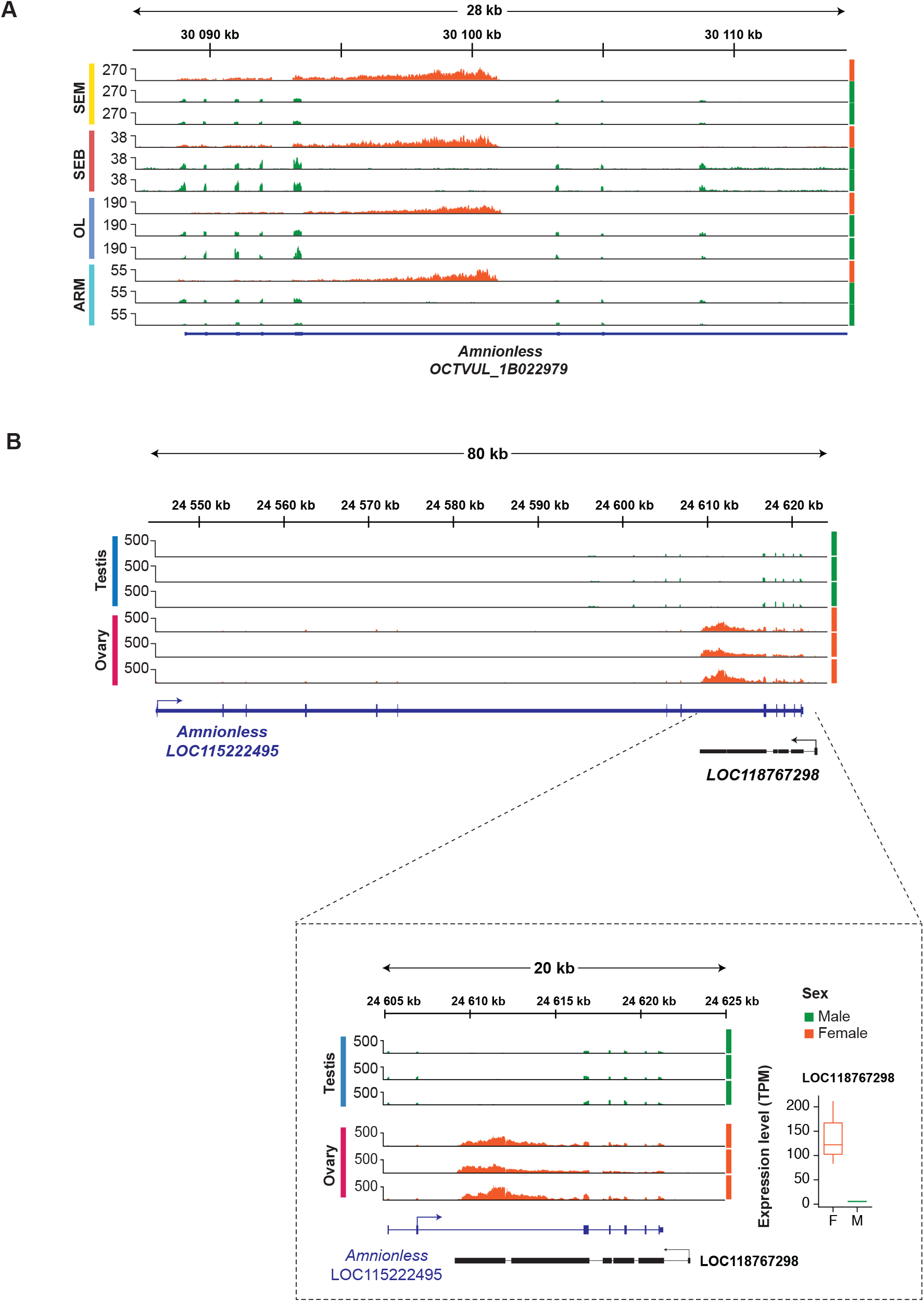
*Zfest* is a female-specific anti-sense lncRNA produced from the Amnionless locus. **A**. Genomic coverage tracks of RNA-seq reads based on *O. vulgaris* tissue RNA-seq data over the Amnionless locus (*OCTVUL_1B022979*) shown for female (in orange) and male (in green) samples. Genomic location of the locus on chromosome Z shown on top. **B**. Top: Genomic coverage tracks of RNA-seq reads based on *O. sinensis* gonadal tissue RNA-seq data over the *Amnionless* (*LOC115222495*) and *Zfest* (*LOC118767298*) loci, for male (in green) and female (in orange) samples. Bottom: Genomic coverage tracks based on the same RNA-seq gonadal tissue data shown for *Zfest* (*LOC118767298*) only, in closer detail. Boxplot showing *Zfest* expression level in TPM for female and male samples. Data shown as median (center line), first and third quartiles (box limits) and 1.5x interquartile range (whiskers).

